# The sources of sex differences in aging in annual fishes

**DOI:** 10.1101/2020.08.25.266411

**Authors:** Martin Reichard, Radim Blažek, Jakub Žák, Petr Kačer, Oldřich Tomášek, Tomáš Albrecht, Alessandro Cellerino, Matej Polačik

## Abstract

Sex differences in lifespan and aging are widespread among animals, with males usually the shorter-lived sex. Despite extensive research interest, it is unclear how lifespan differences between the sexes are modulated by genetic, environmental and social factors. We combined comparative data from natural populations of annual killifishes with experimental results on replicated captive populations, showing that females consistently outlived males in the wild. This sex-specific survival difference persisted in social environment only in two most aggressive species, and ceased completely when social and physical contacts were prevented. Demographically, neither an earlier start nor faster rate of aging accounted for shorter male lifespans, but increased baseline mortality and the lack of mortality deceleration in the oldest age shortened male lifespan. The sexes did not differ in any measure of functional aging we recorded. Overall, we demonstrate that sex differences in lifespan and aging may be ameliorated by modulating social and environmental conditions.

## Introduction

Males and females differ in many demographic and life history parameters, with important consequences for species ecology, evolution, and physiology, as well as for practical and societal outcomes (Trivers 1972, Austad 2006, Regan and Partridge 2013). Inter-sexual differences in lifespan (age at death) and aging (increase in mortality risk associated with deterioration in bodily functions) are widespread among animals, from nematodes to humans (Austad and Fischer 2016). Males are usually the shorter-lived sex (Promislow 2003; Liker and Szekély 2005, Lemaitre et al. 2020), but there is substantial unexplained variation among species and populations (Austad and Fischer 2016). Inter-sexual differences in lifespan and aging appear modulated by environmental and social factors (Austad 2006, Lemaitre et al. 2020), but their effects remain opaque.

Why do males typically express a truncated lifespan in comparison with females? One set of explanations posits that the primary difference stems from genetic and genomic differences between the sexes (Gemmel et al. 2004; Maklakov and Lummaa 2013). In mammals, fruit flies and many other taxa, males are the heterogametic sex and hemizygosity of key genes located on the sex chromosomes resulting in an inability to compensate for the effects of deleterious mutations have been implicated in shorter male lifespan (Trivers 1985; Xirocostas et al. 2020). However, males also show shorter lifespans in many birds and butterflies, despite female heterogamy in those groups (Gotthard et al. 2000; Tompkins and Anderson 2019; Sielezniew et al. 2020). Asymmetry in the inheritance of mitochondria, leading to suboptimal compatibility between mitochondrial and nuclear genomes in males (Frank and Hurst 1996), could explain male-biased mortality and aging in heterogametic taxa (Gemmel et al. 2004). Yet these explanations cannot account for the large variation in the sex bias in lifespan and aging rate seen within species and among inbred laboratory strains housed under contrasting conditions (Austad and Fisher 2016), strongly implicating other factors in driving sex-biased mortality.

Males and females differ in their routes to reproductive success (Trivers 1972). These divergent trajectories arise as a consequence of gamete size disparity which leads to variation in reproductive roles of males and females. This disparity is best explained by sexual selection, with asymmetric variation in reproductive success between the sexes (Andersson 1994). Male reproductive success is often skewed towards a few highly successful individuals while female reproductive success is far less variable (Arnold 1994). Mating system is a key modulator of this variation. In a monogamous mating system, differences in the variance in reproductive success between males and females are trivial. In highly polygynous mating systems, such as those with male harems, a single male may monopolize a large number of females generating highly skewed male reproductive success (Clutton-Brock and Isvaran 2007).

Sexual selection is associated with elevated mortality in the more competitive sex (Székely et al. 2014). Conspicuous signaling to rivals and potential partners directly increases the risk of mortality from predators (Tuttle and Ryan 1991). Male-male competition is also risky and may lead to increased mortality (Beirne et al. 2015). Higher male mortality is often precipitated via alterations to hormonal profiles, resulting in chronic stress (Keller et al. 1992) or elevated testosterone levels (Foo et al. 2017) thereby making individuals more susceptible to infections or physiological deterioration (Moore and Wilson 2002, Gupta et al. 2020).

African annual fishes from the genus *Nothobranchius* are an ideally suited model taxon for biomedical and evolutionary questions related to aging (Cellerino et al. 2016, Hu and Brunet 2018, Cui et al. 2019). Inhabiting ephemeral savanna pools, they have evolved naturally short lifespans which recapitulate typical features of vertebrate aging, including multifarious functional deterioration in old age (Cellerino et al. 2016; Hu and Brunet 2018). In the wild, killifish hatch at the onset of the rainy season from desiccation-resistant eggs. Both sexes grow rapidly and achieve sexual maturity in as few as two weeks (Vrtílek et al. 2018a). Males compete for access to females, with a marked variability in the strength of intra-sexual competition among species (Wildekamp 2004, Genade 2005, Polačik and Reichard 2011; Cellerino et al. 2016). Natural lifespan is limited by desiccation of their habitat, but most fish succumb long before their natal pool desiccates (Vrtílek et al. 2018b). Strikingly, a short lifespan of several months is retained in captivity, where fish are shielded from extrinsic mortality, with captive fish suffering a range of functional declines (Cellerino et al. 2016). In all *Nothobranchius* species for which information on sex chromosomes is available, males are the heterogametic sex, though sex chromosomes are rarely morphologically distinguishable (Krysanov and Demidova 2018).

We combined data from wild populations with experimental results from captive fish to disentangle the causes of differences in lifespan and aging between male and female African annual killifish. Using a set of four species (each replicated as two independent populations), we compared demographic and functional aging between the sexes. Overall, we demonstrate that sex differences in lifespan and aging are primarily modulated by social and environmental conditions.

## Results

### Sex ratio in wild and captive populations

Using adult sex ratios from 376 wild populations (15,968 fish), we found that natural killifish populations in three study species were sex-biased, with significantly more females. Sex ratios in one species (*N. kadleci*) were equal (Fig. 1). This finding corroborated the outcomes of a previous study (Reichard et al. 2014), which was reinforced here using a larger dataset.

**Fig. 1.**
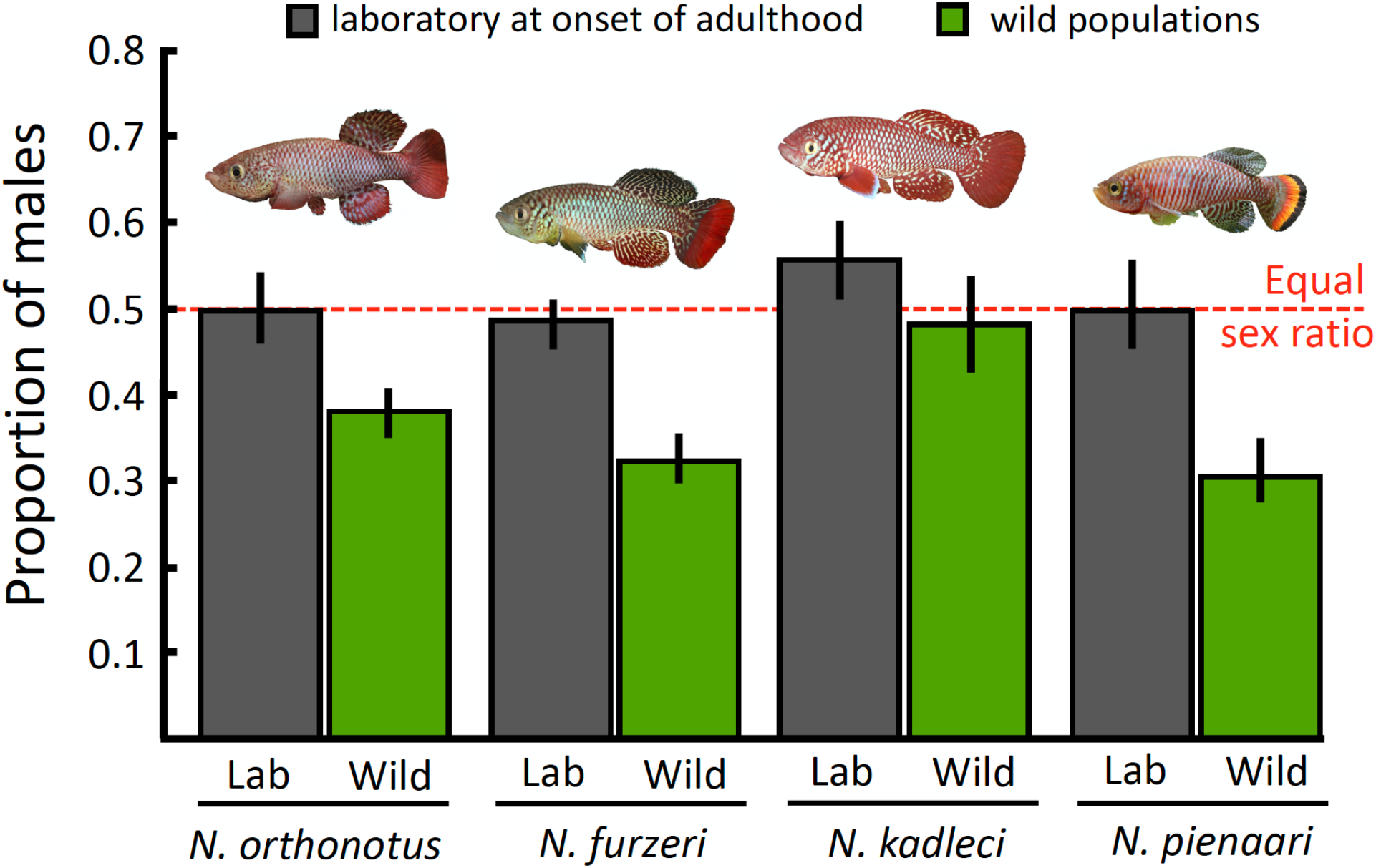
Proportion of males at the onset of sexual maturity in wild-derived laboratory populations (grey columns) and in wild populations (green columns). Means with 95% confidence intervals, back-calculated from outcomes of binomial models, are shown.

Sex ratios in natural populations were recorded throughout the adult phase of life. These results could result from the cumulative effects of biases in primary sex ratios and sex-dependent mortality and to quantify sex-specific survival, an estimate of sex ratios at the onset of adulthood is needed. To obtain these data we raised 63 cohorts of outbred, wild-derived captive populations from study species in protected laboratory conditions. We found that sex ratios in protected conditions were equal in the three study species that exhibited female-biased sex ratio in the wild, while the sex ratio was male-biased in *N. kadleci* – the species with an equal sex ratio in the wild (Fig. 1). This finding implies that mortality of adult males in natural populations was consistently higher than female mortality in all four species.

### Sex differences in lifespan – social and environmental effects

To investigate proximate causes of sex-biased mortality, we raised a set of killifish cohorts from a total of 8 wild-derived populations from all 4 species (Supplementary Table 1) in the laboratory and compared sex differences in lifespan and aging in two contrasting social treatments. By using captive breeding we excluded predation (and predation risk), which has been implicated in sex-biased mortality in wild fish and other animals (Székely et al. 2014), from both treatments. The first treatment comprised replicated social groups of 10-12 fish (equal sex ratio) in which males and females interacted freely, competed and formed dominance hierarchies (N = 84 groups). The second treatment comprised singly-housed fish (N = 178 fish). We predicted that in a captive setting sex differences in mortality would be removed if predation is the source of male-biased mortality. If social stress elevates male mortality, we predicted persistence of male-biased mortality in the social treatment but its disappearance in singly-housed fish treatment. Finally, if intrinsic, sex-specific functional deterioration causes male-biased mortality, we predicted male-biased mortality to persists in both captive treatments.

**Table 1.** Sex-specific median lifespan estimates (with 95% confidence intervals estimated from the *survfit* function) in the social and single housing treatments. NAs represent cases where the upper confidence interval cannot be reliably calculated.

| Median lifespan (95% CI) |  |  |  |
| --- | --- | --- | --- |
| Species | Sex | Social | Single |
| *N. orthonotus* | Females | 256 (214-289) | 232 (193-352) |
|  | Males | 180 (134-197) | 271 (215-368) |
| *N. furzeri* | Females | 150 (131-176) | 121 (104-NA) |
|  | Males | 121 (115-134) | 117 (111-145) |
| *N. kadleci* | Females | 172 (148-272) | 166 (103-240) |
|  | Males | 178 (161-206) | 113 (97-NA) |
| *N. pienaari* | Females | 282 (221-338) | 314 (242-347) |
|  | Males | 251 (227-286) | 297 (235-446) |

We found support for predation-related and social stress-related decreases in male lifespan. First, sex differences in lifespan in social tanks persisted in two species – *N. orthonotus* (z = 4.84, P < 0.001; with male median lifespan 42%, i.e. 76 days shorter) and *N. furzeri* (z = 2.64, P = 0.008; male lifespan 24%, i.e. 29 days shorter) but disappeared in the other two – *N. kadleci* (z = 0.24, P = 0.81) and *N. pienaari* (z = 0.26, P = 0.79; Table 1). This demonstrates that the absence of predation eliminated the sex bias in mortality in two species, but not in the other two. This interspecific variation in socially-induced sex bias in lifespan tightly covaries with the level of male aggressiveness, which is markedly higher in *N. orthonotus*, followed by *N. furzeri, N. kadleci* and *N. pienaari* (Wildekamp 2004, Genade 2005, Polačik and Reichard 2011).

Second, there was no sex bias in lifespan when fish were housed singly (*N. orthonotus*: z = 0.14, P = 0.89; *N. furzeri*: z = 0.50, P = 0.62; *N. kadleci*: z = 1.28, P = 0.20; *N. pienaari*: z = 1.54, P = 0.12). Lifespan estimates of fish that lived in social and singly-housed treatments were congruent (95% confidence intervals for median lifespans overlapped) except for an increase in median lifespan in singly-housed *N. orthonotus* males (Table 1), the most aggressive species. This finding implies that male-male aggression considerably decreased male lifespan in *N. orthonotus* in a social setting.

Finally, we tested the hypothesis that the sex bias in mortality observed in challenging natural conditions disappeared in more benign conditions in captivity, using one population of *N. furzeri*. We replicated a strong circadian fluctuation in water temperature (from 20±1°C in early morning to 35±1°C in late afternoon), which is characteristic of the natural environment (Žák et al. 2018) and exceeds killifish preferred temperature variation by 6°C (Polačik et al. 2016, Žák et al. 2018). This thermal challenge is not being employed in standard breeding protocol for captive killifish (Polačik et al. 2016), which we imposed in our main experiment. We found that even under these challenging environmental conditions, there was no sex bias in mortality in singly-housed *N. furzeri* (thermally fluctuating environment: z = 0.55, P = 0.582, N = 45 *N. furzeri* kept as singly-housed fish; control stable temperature of 27.5±1 °C: z = 1.04, P = 0.297, N = 45).

### Sex differences in actuarial aging

Sex-biased lifespan can arise from differences in baseline mortality (i.e. one sex experiencing persistently higher mortality) or demographic rate of aging (i.e. a steeper increase in mortality with age in one sex). These two sources of differential mortality can be best estimated with a Gompertz model of increasing failure time (Bronikowski et al. 2011; Boonekamp et al. 2020). We fitted a set of models (Colchero et al. 2012) to our data from the social treatment, in which shorter male lifespan was detected, and confirmed that Gompertz-family models well approximated observed mortality patterns (Supplementary Table 2). We used Bayesian survival trajectory analysis (BaSTA; Colchero et al. 2012) to estimate intersexual differences in actuarial ageing within populations of species with sex-biased mortality.

**Table 2.** The effects of sex, age and their interaction on proliferative changes to liver (a), deposition of lipofuscin (b) and proliferative changes to kidney (c). Parameter estimates and z-statistics of Generalized Mixed Models are presented.

| Organ impairment | Factor | Estimate | ± s.e. | z-value | P |
| --- | --- | --- | --- | --- | --- |
| Proliferative changes - liver | Intercept | 0.761 | ± 0.242 | 3.14 | 0.002 |
| | Sex(males) | -0.110 | ± 0.218 | -0.50 | 0.615 |
| | Age(young) | -0.240 | ± 0.225 | 1.07 | 0.287 |
| | Sex x Age | 0.449 | ± 0.310 | 1.45 | 0.147 |
| Lipofuscin - liver | Intercept | 3.099 | ± 0.263 | 11.78 | <0.001 |
| | Sex(males) | -0.045 | ± 0.259 | -0.17 | 0.862 |
| | Age(young) | 0.071 | ± 0.045 | 1.57 | 0.115 |
| | Sex x Age | -0.195 | ± 0.299 | -0.65 | 0.514 |
| Proliferative changes - kidney | Intercept | 0.338 | ± 0.347 | 0.97 | 0.331 |
| | Sex(males) | -0.481 | ± 0.281 | -1.71 | 0.087 |
| | Age(young) | -0.637 | ± 0.353 | -1.81 | 0.071 |
| | Sex x Age | 0.551 | ± 0.474 | 1.16 | 0.247 |

In *N. orthonotus*, the species with the greatest contrast in lifespan between the sexes, male-biased mortality was affected by stronger baseline mortality in males and not their higher rate of aging, consistently across both study populations (Figure 2a, b). In *N. furzeri*, the second species with male-biased mortality in the social treatment, a Gompertz-logistic model (which includes an additional parameter describing deceleration in the aging rate in old age) gave a significantly better fit to observed data than a simple Gompertz model (Supplementary Table 2). We detected lower mortality deceleration in old age (parameter *c*) in one *N. furzeri* population (Figure 2c) and no intersexual difference in Gompertz model parameters in the second population (Figure 2d).

**Fig. 2.**
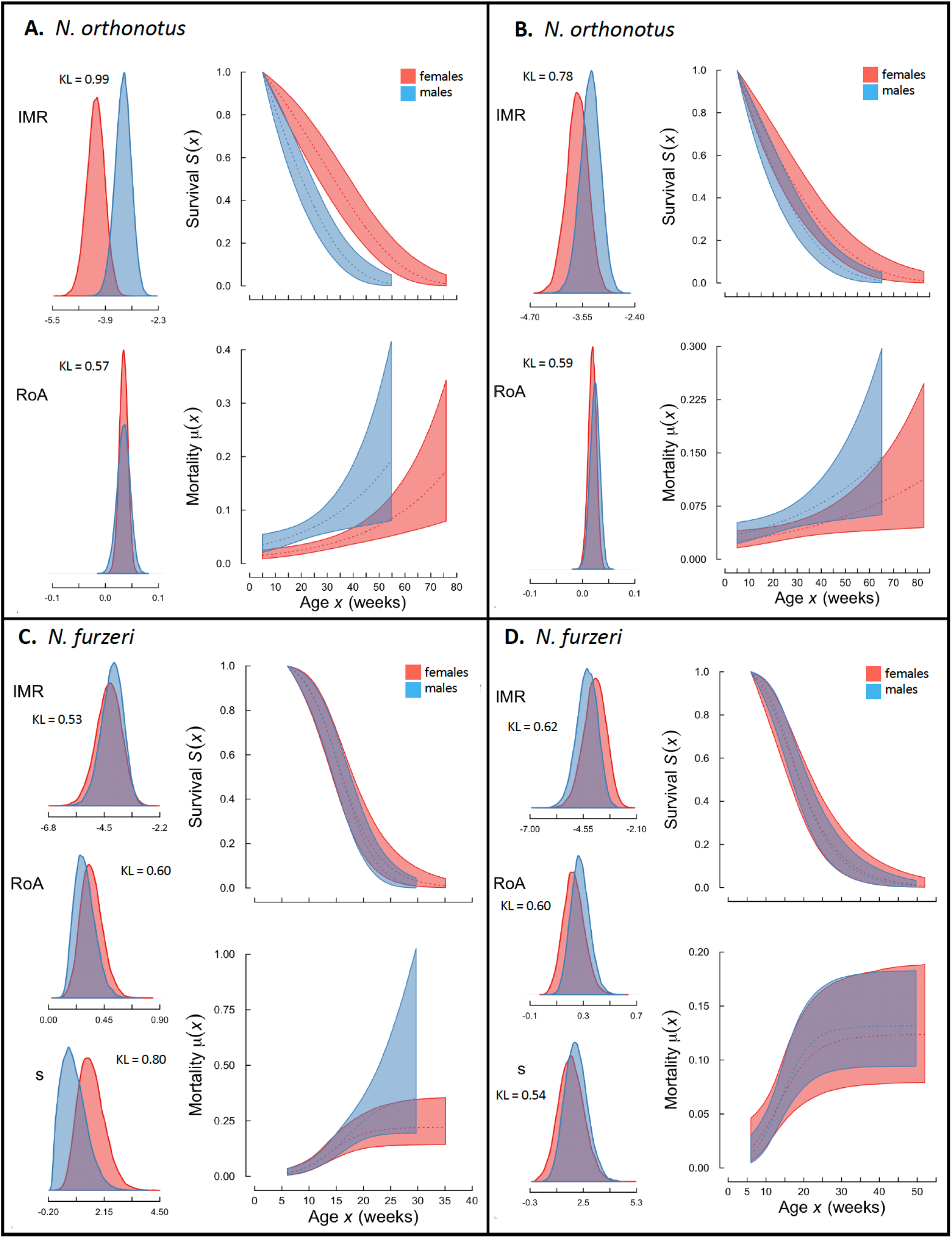
Sex-specific posterior distribution of baseline mortality (IMR), rate of aging (RoA), survival, and mortality risk estimated from Gompertz model for both *N. orthonotus* populations (A, B) and both *N. furzeri* populations (C, D) in the social treatment. In *N. furzeri*, the posterior distribution for aging deceleration parameter (*s*) is also presented. KL denotes Kullback-Leibler discrepancy criterion, with values >0.8 considered as a substantial difference.

### Sex differences in functional aging

In a protected environment, mortality derives from deterioration in bodily function. We contrasted data on biomarkers of cellular and physiological aging between males and females kept in the social treatment. We analyzed markers of oxidative stress to lipids, proteins and DNA in liver, brain and heart tissues of young (14 weeks) and old (24 weeks) fish from all 8 experimental populations using liquid chromatography-electrospray-high resolution mass spectrometry. A PCA-based composite value well approximated oxidative stress across tissues and markers (PC1 explained 72% of variation, Supplementary Table 3). Oxidative stress increased with age (LMM: t_140_ = 24.89, P <0.001) but did not differ between the sexes in either absolute values (t_140_ = 0.14, P = 0.886) nor in the steepness of its increase with age (sex by age interaction: t_140_ = 0.49, P = 0.629). The same outcome was obtained from species-specific analyses (Supplementary Table 4) and for biomarker- and tissue-specific analyses (Supplementary Table 5). Hence, there was no detectable intersexual difference in oxidative stress.

Using a different set of individuals, we compared the deposition of lipofuscin in liver tissue, as a biomarker of cellular aging. Lipofuscin is an aggregate of oxidized proteins that accumulates in aged post-mitotic cells (Jung et al. 2006). Lipofuscin deposition did not differ between males and females (LMM with Poisson error distribution: z = 0.17, P = 0.862, N = 75 fish) and we found no significant increase in lipofuscin accumulation with age (z = 1.57, P = 0.115; sex by age interaction: z = 0.65, P = 0.514). Finally, as *Nothobranchius* killifish age they suffer proliferative changes leading to organ dysfunction that is linked to mortality and which can be revealed by histopathological examination (di Cicco et al. 2010, Baumgart et al. 2015). We found no intersexual differences in proliferative changes in the kidney (P = 0.847) or liver (P = 0.115; Table 2).

## Discussion

Intersexual differences in lifespan and aging are widespread among taxa, but despite a substantial research interest, it is still not clear how genetic differences between the sexes are modulated by environmental and social factors (Austad 2006; Gordon et al. 2017; Lemaitre et al. 2020). Using eight populations from four closely related annual killifish species, we combined comparative and experimental approaches to demonstrate that female bias in wild annual killifish populations arises from a combination of higher extrinsic male mortality in natural populations and higher intrinsic mortality linked to social interactions, rather than from generalized intersexual differences in functional deterioration. Females consistently outlived males in the wild, but this difference persisted in social tanks only in more aggressive species, and ceased when fish were housed singly. Increased baseline mortality, not an earlier or faster rate of aging was primarily responsible for a shorter male lifespan in a social setting. Importantly, there were no differences between the sexes in a series of measures of functional aging (oxidative stress, lipofuscin deposition, or age-related proliferative changes in liver and kidney).

The impacts of sexual selection explained male-biased mortality in natural and experimental annual killifish populations. In the wild, there is evidence that males suffer elevated predation. Annual killifish are highly sexually dichromatic, with brightly colored males and dull females (Sedláček et al. 2014), and visually hunting birds, such as herons and kingfishers, are the main predators of annual killifish (Haas, 1976, Keppeler et al. 2016; Reichard and Polačik 2019). A mortality cost of showy sexually-selected traits is a well-recognized source of intersexual differences in lifespan (Promislow et al. 1992; Székely et al. 2014, Lemaitre et al. 2020). In addition, we found male-male competition for mating opportunities significantly contributed to elevated male mortality in more aggressive species. Notably, we observed male combat-related injuries in five *N. orthonotus* and three *N. furzeri* males that died in the social group treatment. Persistent stress (Keller et al. 1992), possibly mediated by elevated levels of corticosteroids (Foo et al. 2017), is frequently associated with increased mortality (Moore and Wilson 2002), but hormonal profiles were not measured in our study. Unexpectedly, we detected no functional characteristics underlying a higher male baseline mortality in our study, despite using measures of physiological aging that were previously found to be suitable biomarkers of functional decline as they predictably varied with age and among killifish species and populations (Terzibasi-Tozzini et al. 2013; Baumgart et al. 2015; Blažek et al. 2017). This is comparable to well-described male-female health-survival paradox in humans, where woman outlive men despite experiencing greater levels of functional problems at older age (reviewed in Gordon et al. 2017).

One species (*N. kadleci*) exhibited equal adult sex ratios in the wild and a male-biased sex ratio at sexual maturity in captivity, consistently across populations and cohorts. Sowersby et al. (2020) hypothesized that sex ratios can evolve extremely rapidly in killifish, demonstrating large interspecific differences among adult sex ratios across 15 annual and non-annual killifish species raised in the lab. We propose that male-biased sex ratio in *N. kadleci* might have evolved as compensatory mechanism to mitigate male-biased mortality in natural populations. Some wild populations in our dataset presented extremely female-skewed sex ratios (e.g. in one natural population we have collected one male and 44 females), and production of male-biased progeny would be adaptive in such populations in accordance with Fisher’s principle (Fisher 1930). Ongoing cyto(genetic) research aims to characterize the nature of this potential compensatory mechanism.

Despite an enormous research effort, a comprehensive causal understanding of sex differences in lifespan and aging remains elusive, probably because it comprises a series of complex underlying sources. Mammals are arguably the best studied vertebrate taxon with respect to aging. A recent comparative study that combined data from 101 mammalian species demonstrated that females lived on average 19% longer than conspecific males but without finding any consistent intersexual differences in aging rates (Lemaitre et al. 2020), in line with our experimental results with killifishes. The fact that heterospecific-sex disadvantage is much stronger in male heterogametic systems (21% longer lifespan of homogametic females) than female heterogametic systems (7% longer lifespan of homogametic males) (Xirocostas et al. 2020) highlights the importance of reproductive roles and mating systems in shaping intersexual lifespan differences. Here, we have demonstrated that sexual selection, which acts differently on the sexes, substantially alters sex differences in lifespan and aging through multiple processes even within an ecologically and evolutionary discrete lineage, and that these effects are strongly moderated by the social and environmental setting.

## Materials and Methods

### Sex ratio estimates from wild populations

We estimated the sex ratio in wild populations of all four study species from 10 field trips to Mozambique conducted between 2008 and 2015. *Nothobranchius* populations contain a single age cohort since fish hatch in synchrony soon after rains fill their natal pools with water (Vrtílek et al. 2018b). The age of fish when sex ratios were estimated was unknown. At each site, fish were collected using a triangular dip net (45 x 45 cm, mesh size 5 mm) or beach seine (length 2.7 m, depth 0.7 m, mesh size 4 mm). The method retained adult killifish unselectively and there was no sex bias in the probability of capture, confirmed by a combination of capture-mark-recapture studies and removal sampling (Reichard et al. 2014, Vrtílek et al. 2018b). Fish were sorted into species and sexed on the bank, counted and released back to the pool. Details for data collection are provided in Reichard et al. (2014); the new samples used in the present study were collected following an identical protocol. We only used estimates based on at least 6 individuals of a given species in further analyses. Sex ratios were analyzed using a Generalized Linear Mixed Model (GLMM) with binomial error structure (male to female ratio) and log-link function in the *lmer* package (Pinheiro and Bates 2000), where *Species* were treated as fixed factor and *Year* and *Site* as random factors.

Data on sex ratios at the start of sexual maturity in wild-derived captive populations were collected in captivity from 63 cohorts. Within each cohort, fish were hatched on the same day, following standard husbandry protocol (Polačik et al. 2016). The number of males and females was estimated at age when all fish in the cohort were sexually mature (typically 4-5 weeks). Data were analyzed using a Generalized Linear Model (GLM) with binomial error structure and log-link function.

### Experimental populations

We used fish from four related *Nothobranchius* species from southern and central Mozambique (Reichard et al. 2017). For laboratory experiments, each species was represented by two independent populations, originating from separate intraspecific lineages (Bartáková et al. 2015). Experimental fish were F1 descendants of wild parents collected in Mozambique. The locations of source populations are presented in Supplementary Table S1. Eggs of parental fish were stored in an incubator (Pollab, Q-CELL 60-240) at 24±0.5°C for at least 16 weeks following standard husbandry protocols (Polačik et al. 2016). The experiment was divided into two phases for logistic reasons (capacity of experimental facility). Work on *N. furzeri* and *N. kadleci* was completed in September 2011 - December 2012, followed by work on *N. orthonotus* and *N. pienaari* (May 2013 - March 2015).

Experimental fish were hatched simultaneously by watering the incubation substrate with dechlorinated tap water (16°C). From the age of 2-10 days (depending on size of the juveniles, but before the sexes could be separated) fish were housed either in social tanks (24 L) or individually (2L tanks), providing identical fish density between treatments. During the juvenile period, dead experimental fish were replaced with fish of the same age and housing history from stock tanks. At the age of 6-7 weeks, fish in social tanks were marked with a single Visible Implant Elastomer tag (Northwestern Marine Technology) to enable individual recognition, except for *N. pienaari* due to its small size. Previous studies have shown no negative effect of marking on subsequent survival (Sandford et al. 2020). Nine to twelve tanks were used for each study population, with initial density of 12 fish per 24L tank, except for *N. orthonotus* (the largest species) where the density was 10 fish per tank. Water quality was maintained using air-driven filters and 25-30% of water was exchanged every 2-3 days. Individually housed fish were kept in 2L tanks in two separate recirculating systems (Aquamedic, Germany, www.aqua-medic.de), with 45 fish per species (22-23 fish per population). All fish were kept under a 12 h:12 h light:dark regime in aged tap water (conductivity 550 μS.cm^-1^), at a water temperature of 26 ±2 °C. Fish were fed twice each day to satiation during the first month and once a day thereafter. Fish were initially fed with live *Artemia* nauplii and weaned to chopped bloodworm (*Chironomus* larvae) and *Tubifex* from the age of 10-30 days. All tanks received the same ration (approximately 15% of body mass of the fish in the tank). Full details are provided in Blažek et al. (2017).

To test the effect of unfavorable environmental conditions arising from fluctuating temperature (Thomas et al. 1986), we compared sex differences in lifespan in a cohort of individually housed fish (*N. furzeri*, population A) that experienced either a stable temperature (mean ± SD: 27.5 ± 1°C; control fish) or fluctuating temperature (from 20±1°C in early morning to 35±1°C in late afternoon). The limits for the fluctuating temperature reflected the diurnal change in water temperature that killifish typically experience in the wild (Žák et al., 2018). A fluctuating temperature was achieved by a combination of an aquarium chiller (TECO TR 10, Italy, www.tecoonline.com) and three aquarium heaters (2× 200 W and 1× 100 W, Eheim/Jäger, Wüstenrot, Germany). Stable temperature in the control group was regulated with one 100 W heater.

### Lifespan estimates

All tanks were monitored daily for dead fish. Survival was estimated from the age when all fish of a given species were sexually mature (5 weeks in *N. orthonotus*, 6 weeks in *N. furzeri* and *N. kadleci*, 8 weeks in *N. pienaari*). Sex differences in mortality were analyzed using species-specific Mixed Effects Cox Proportional-Hazards Models (*coxme* package) (Therneau 2015a) with *Sex* as a fixed factor and *Population* as a random factor. Note that analysis using *Population* as the fixed factor (*coxph* function in *survival* package, including population by sex interaction) generated an identical interpretation. Analyses were completed separately for each species and social environment. Fish removed from social tanks for the analysis of functional aging were censored in the survival analysis at the age of removal.

### Actuarial aging

Sex-specific mortality hazards were modelled using Bayesian Survival Trajectory Analysis (*BaSTA* package) (Colchero et al. 2012). First, we used the *multibasta* command to test whether Gompertz-family models provided a good fit to population-specific survival data. We fitted three basic models (Weibull, Gompertz, Logistic), each with three shape parameters (simple, Makeham, Siler/bathtub) and then compared the fits using Deviance Information Criterion (DIC), a Bayesian equivalent of Akaike Information Criterion. We used 4 runs of each model, each with 150,000 MCMC iterations, burn in of 15,000 and thinning by sampling every 50th estimate. We analyzed each population separately as we knew a priori that populations within species differ in lifespan (Blažek et al. 2017) and, unlike for survival analysis, they cannot be entered as random effects to BaSTA. Gompertz-family models were chosen to provide the most unambiguous demographic interpretation of the parameters (Bronikowski et al. 2011; Boonekamp et al. 2020) and a good fit to the datasets. The Gompertz model assumes that aging starts at species-specific age, with one parameter (intercept, *Initial mortality rate, IMR*) describing age-independent mortality (baseline mortality) and the second parameter (slope, *Rate of Aging, RoA*) describing the increase in mortality with age (Pletcher et al. 2000). In both *N. furzeri* populations, deceleration in aging was apparent at old age, probably arising from intra-population heterogeneity (Chen et al. 2013), and Gompertz-logistic models were used as their DIC was considerably lower than a simple Gompertz model. A Gompertz-logistic model estimates a third parameter (*s*) which models deceleration of aging rate at old age. The final models were run with 400,000 MCMC iterations, burn in of 50,000 and thinning by sampling every 50th estimate to provide a posterior distribution of parameters for each species. Model parameters were compared between males and females using Kullback-Leibler discrepancy criterion (KL). The KL varies between 0.5 (complete overlap) to 1.0 (no overlap) with the values > 0.8 are considered as a substantial difference (Colchero et al. 2012).

### Oxidative stress

Subsamples of 20 young and old fish from social treatment (young fish: 12 weeks; old fish: 26 weeks in *N. furzeri* and *N. kadleci* and 30 weeks in *N. orthonotus* and *N. pienaari*), were sacrificed for per species (equal representation of sex and populations). Their brain, liver and heart were flash frozen in liquid nitrogen. Tissues were homogenized in acetonitrile with deuterium-labelled internal standards, and oxidation products of nucleic acids (8-hydroxy-2’-deoxyguanosine (8-OHdG); 8-hydroxyguanosine (8-OHG)), proteins (o-Tyrosine, 3-nitrotyrosine, 3-chlorotyrosine) and lipids (8-isoprostane) were determined by liquid chromatography-electrospray-high resolution mass spectrometry (HPLC-ESI-HRMS) as described in Blažek et al. (2017). *N. orthonotus* and *N. pienaari* from the main experiment were used, but new cohorts of *N. furzeri* and *N. kadleci* were raised using identical conditions (Blažek et al. 2017). Within individuals, data from the three organs and biomarkers were collinear. We combined data across organs and biomarkers using Principal Component Analysis (Supplementary Table S3). The first PC explained 71.8% of variability and had positive loadings with all biomarkers across all tissues (0.77-0.92). We used a Linear Mixed Model to test how the sexes differed in oxidative stress (using *PC1* as a fixed effect) and how increase in oxidative stress with age varied between the sexes (i.e. a *Sex* by *Age* interaction). *Population ID* was a random effect. In addition to an overall test of oxidative stress, we used organ-specific and biomarker-specific analyses (Supplementary Tables S4 and S5), which were fully concordant with the PC1-based analysis.

### Histopathology

Another sample of young (age 14 weeks) and old (age 23 weeks) *N. furzeri* and *N. kadleci* from social treatment was sacrificed (20 fish per age and species). Liver and kidney were preserved in Baker’s solution, embedded in Paraplast, sectioned (5 μm) and stained in H&E. From the histological slides, the incidence of proliferative changes was scored using a 5-grade pathological scale (Di Cicco et al. 2011) (score 0-4, 0: no proliferation, 4: >50% of tissue filled with proliferative cells). Data were analyzed using Cumulative Link Mixed models for ordinal data in the package *ordinal* (Christensen, 2019), with *Sex, Age* and their interaction as fixed factors, and *Population ID* as a random factor. The amount of lipofuscin particles was estimated from separate slides (unstained sections) using a Leica confocal fluorescent microscope. Nine slides were analyzed for each individual. Excitation wavelength was set to 488 nm (confocal parameters as pinhole, photo-multiplier and laser intensity were fixed). Images were imported to *imageJ* and the number of lipofuscin (fluorescing) particles counted. Data were analyzed using Generalized Linear Mixed models with Poisson errors (counts), with *Sex, Age* and their interaction as fixed factors, and *Population ID* and *Individual ID* as random effects.

### Data availability

All data associated with paper are available in the Figshare repository (doi: 10.6084/m9.figshare.12752648).

## ACKNOWLEDGEMENTS

Funding came from Czech Science Foundation (19-01781S) to M.R. All work was carried out in accordance with relevant guidelines and regulations. Fieldwork complied with legal regulations of Mozambique (collection licenses: DPPM/053/7.10/08, 175/154/ IIP/2009/DARPE, DPPM/083/7.10/10, DPPM/330/7.10/10, DPPM/069/7.10/11, DPPM/088/7.10/12). Experimental work was approved by the Ethical Committee of the Institute of Vertebrate Biology (No. 163-12) and by Ministry of Agriculture (CZ 62760203) in accordance with legal regulations of the Czech Republic. We thank C. Smith for valuable comments.

## AUTHOR CONTRIBUTIONS

MR conceived and designed the study, conducted statistical analyses and drafted the manuscript. MR, RB and MP collected data on wild populations. RB completed the experiment with captive fish, with the assistance of MP. JZ collected data from fluctuating temperature. PK analyzed tissues for oxidative stress. OT designed and interpreted oxidative stress data. TA established and managed team for oxidative stress analysis. AC and RB performed histological analysis. All authors contributed to the final text.

