## Supplemental Data for "The sources of sex differences in aging in annual fishes"

**Table S1.** Overview of study populations, with sample size for social tank treatment, collection code for each population and GPS coordinates of the source habitat and its altitudinal position.

| Population | $N_S$ | Collection code MZCS | GPS coordinates | Altitude (m) |
|---|---|---|---|---|
| *N. orthonotus* A | 131 | NO2 | S24.064 E32.732 | 79 |
| *N. orthonotus* B | 139 | NO528 | S19.697 E34.783 | 38 |
| *N. furzeri* A | 102 | NF222 | S21.873 E32.801 | 158 |
| *N. furzeri* B | 99 | NF121 | S24.358 E32.974 | 30 |
| *N. kadleci* A | 108 | NK91 | S20.688 E34.106 | 64 |
| *N. kadleci* B | 101 | NK430 | S19.281 E34.221 | 52 |
| *N. pienaari* A | 140 | NP505 | S23.530 E32.578 | 129 |
| *N. pienaari* B | 157 | NP514 | S19.700 E34.785 | 36 |

**Table S2.** Relative fit of demographic models of aging in BaSTA ranked on the basis of difference in DIC (Deviance Information Criterion). Models used in species-specific analyses are indicated in bold. Note that *N. kadleci* and *N. pienaari* did not show any difference in lifespans between males and females in a captive setting, therefore sex-specific aging patterns were not compared.

| Population | Rank | Model | Shape | ΔDIC |
|---|---|---|---|---|
| *N. orthonotus* | 1 | **Gompertz** | **simple** | **0.0** |
| *Dry* | 2 | Weibull | simple | 0.9 |
| | 3 | Weibull | Makeham | 1.1 |
| | 4 | Gompertz | Makeham | 2.1 |
| | 5 | Logistic | Makeham | 2.2 |
| | 6 | Logistic | simple | 3.0 |
| | 7 | Weibull | bathtub | 6.3 |
| | 8 | Gompertz | bathtub | 6.4 |
| | 9 | Logistic | bathtub | 7.7 |
| *N. orthonotus* | 1 | Weibull | simple | 0.0 |
| Wet | 2 | **Gompertz** | **simple** | **2.8** |
| | 3 | Logistic | simple | 3.0 |
| | 4 | Weibull | Makeham | 3.0 |
| | 5 | Logistic | Makeham | 3.3 |
| | 6 | Weibull | bathtub | 8.5 |
| | 7 | Gompertz | Makeham | 9.1 |
| | 8 | Logistic | bathtub | 9.4 |
| | 9 | Gompertz | bathtub | 10.2 |
| *N. furzeri* | 1 | **Logistic** | **simple** | **0.0** |
| *Dry* | 2 | Logistic | Makeham | 3.0 |
| | 3 | Weibull | simple | 4.6 |
| | 4 | Logistic | bathtub | 7.1 |
| | 5 | Weibull | Makeham | 7.9 |
| | 6 | Weibull | bathtub | 11.7 |
| | 7 | Gompertz | simple | 27.9 |
| | 8 | Gompertz | Makeham | 32.6 |
| | 9 | Gompertz | bathtub | 38.5 |
| *N. furzeri* | 1 | **Logistic** | **simple** | **0.0** |
| Wet | 2 | Weibull | simple | 1.7 |
| | 3 | Logistic | Makeham | 4.5 |
| | 4 | Weibull | Makeham | 6.4 |
| | 5 | Logistic | bathtub | 7.6 |
| | 6 | Weibull | bathtub | 12.1 |
| | 7 | Gompertz | simple | 19.8 |
| | 8 | Gompertz | Makeham | 27.2 |
| | 9 | Gompertz | bathtub | 29.2 |

**Table S3.** Principal Component Analysis (PCA) combining oxidative stress in liver, brain and heart using biomarkers of damage to lipids, proteins and DNA.

| Organ | Biomarker | Oxidative damage to: | PC 1 loading | PC 2 loading | PC 3 loading | PC 4 loading | PC 5 loading |
|---|---|---|---|---|---|---|---|
| Liver | 8-isoprostane | lipids | 0.828 | 0.543 | 0.056 | 0.001 | 0.001 |
| Liver | o-tyrosine | proteins | 0.765 | 0.561 | 0.105 | 0.087 | -0.060 |
| Liver | 3-nitrotyrosine | proteins | 0.779 | 0.541 | 0.076 | -0.023 | 0.141 |
| Liver | 3-chlorotyrosine | proteins | 0.807 | 0.543 | 0.071 | -0.026 | -0.039 |
| Liver | 8-OHdG | DNA | 0.813 | 0.519 | 0.073 | 0.005 | 0.045 |
| Liver | 8-OHG | DNA | 0.792 | 0.570 | 0.071 | 0.003 | -0.077 |
| Brain | 8-isoprostane | lipids | 0.886 | -0.331 | 0.299 | -0.003 | -0.001 |
| Brain | o-tyrosine | proteins | 0.855 | -0.348 | 0.272 | -0.005 | -0.149 |
| Brain | 3-nitrotyrosine | proteins | 0.867 | -0.330 | 0.302 | -0.016 | -0.049 |
| Brain | 3-chlorotyrosine | proteins | 0.861 | -0.340 | 0.294 | -0.001 | -0.018 |
| Brain | 8-OHdG | DNA | 0.829 | -0.347 | 0.313 | 0.064 | 0.253 |
| Brain | 8-OHG | DNA | 0.869 | -0.342 | 0.278 | -0.014 | -0.047 |
| Heart | 8-isoprostane | lipids | 0.923 | -0.129 | -0.305 | -0.062 | -0.049 |
| Heart | o-tyrosine | proteins | 0.894 | -0.158 | -0.385 | 0.115 | -0.012 |
| Heart | 3-nitrotyrosine | proteins | 0.895 | -0.142 | -0.339 | -0.220 | 0.006 |
| Heart | 3-chlorotyrosine | proteins | 0.856 | -0.167 | -0.419 | 0.179 | 0.005 |
| Heart | 8-OHdG | DNA | 0.873 | -0.154 | -0.323 | -0.296 | 0.050 |
| Heart | 8-OHG | DNA | 0.848 | -0.229 | -0.354 | 0.240 | 0.012 |

| **Principal components** | | | | | |
|---|---|---|---|---|---|
| Eigenvalue | 12.93 | 2.65 | 1.31 | 0.26 | 0.13 |
| Variance explained (%) | 71.8 | 14.7 | 7.3 | 1.4 | 0.7 |
| Cumulative variance explained (%) | 71.8 | 86.6 | 93.8 | 95.3 | 96.0 |

**Table S4**. The fixed effect parameter estimates of Generalized Mixed Models for species-specific oxidative stress on combined data across tissues and biomarkers (PC1).

| Species | Factor | Estimate | ± s.e. | t-value | P |
|---|---|---|---|---|---|
| *N. orthonotus* | Intercept | -3.00 | ± 1.56 | -1.92 | 0.284 |
| | Sex(males) | 0.39 | ± 0.61 | 0.64 | 0.528 |
| | Age(old) | 6.87 | ± 0.55 | 12.41 | <0.001 |
| | Sex x Age | -0.15 | ± 0.93 | -0.14 | 0.894 |
| *N. furzeri* | Intercept | -2.52 | ± 1.32 | -1.91 | 0.295 |
| | Sex(males) | -0.45 | ± 0.44 | -1.01 | 0.320 |
| | Age(old) | 5.61 | ± 0.46 | 12.29 | <0.001 |
| | Sex x Age | 0.96 | ± 0.66 | 1.45 | 0.158 |
| *N. kadleci* | Intercept | -2.57 | ± 0.53 | -4.87 | 0.051 |
| | Sex(males) | -0.15 | ± 0.44 | -0.34 | 0.737 |
| | Age(old) | 6.10 | ± 0.44 | 13.99 | <0.001 |
| | Sex x Age | 1.06 | ± 0.74 | 1.42 | 0.164 |
| *N. pienaari* | Intercept | -3.61 | ± 1.58 | -2.29 | 0.243 |
| | Sex(males) | 0.08 | ± 0.57 | 0.16 | 0.886 |
| | Age(old) | 6.37 | ± 0.57 | 11.27 | <0.001 |
| | Sex x Age | -0.44 | ± 0.80 | -0.55 | 0.584 |

**Table S5**. The fixed effect parameter estimates of Generalized Mixed Models for tissue-specific oxidative stress for specific biomarkers (lipids: 8-isoprostane, proteins: o-tyrosine, nucleic acids: (8-hydroxy-2'-deoxyguanosine, 8-OHdG).

**(a) Liver tissue**

| Species | Factor | Estimate | ± s.e. | t-value | P |
|---|---|---|---|---|---|
| 8-isoprostane | Intercept | 1.33 | ± 0.06 | 20.59 | <0.001 |
| | Sex(males) | -0.01 | ± 0.04 | -0.42 | 0.676 |
| | Age(old) | 0.48 | ± 0.04 | 11.35 | <0.001 |
| | Sex x Age | 0.09 | ± 0.07 | 1.34 | 0.183 |
| o-tyrosine | Intercept | 1.31 | ± 0.07 | 19.51 | <0.001 |
| | Sex(males) | -0.04 | ± 0.05 | -0.78 | 0.435 |
| | Age(old) | 0.49 | ± 0.05 | 9.75 | <0.001 |
| | Sex x Age | 0.04 | ± 0.08 | 0.46 | 0.647 |
| 8-OHdG | Intercept | 0.38 | ± 0.02 | 19.06 | <0.001 |
| | Sex(males) | -0.01 | ± 0.01 | -0.61 | 0.541 |
| | Age(old) | 0.15 | ± 0.01 | 10.62 | <0.001 |
| | Sex x Age | 0.01 | ± 0.02 | 0.67 | 0.501 |

**(b) Heart tissue**

| Species | Factor | Estimate | ± s.e. | t-value | P |
|---|---|---|---|---|---|
| 8-isoprostane | Intercept | 7.93 | ± 0.60 | 13.23 | <0.001 |
| | Sex(males) | -0.22 | ± 0.31 | -0.72 | 0.472 |
| | Age(old) | 5.63 | ± 0.31 | 18.52 | <0.001 |
| | Sex x Age | 0.33 | ± 0.16 | 0.71 | 0.478 |
| o-tyrosine | Intercept | 24.18 | ± 1.92 | 12.63 | <0.001 |
| | Sex(males) | -1.06 | ± 1.11 | -0.96 | 0.339 |
| | Age(old) | 16.67 | ± 1.10 | 15.19 | <0.001 |
| | Sex x Age | 1.08 | ± 1.68 | 0.64 | 0.522 |
| 8-OHdG | Intercept | 6.97 | ± 0.52 | 13.38 | <0.001 |
| | Sex(males) | -0.10 | ± 0.36 | -0.29 | 0.774 |
| | Age(old) | 5.28 | ± 0.35 | 14.91 | <0.001 |
| | Sex x Age | -0.40 | ± 0.54 | -0.74 | 0.459 |

**(c) Brain tissue**

| Species | Factor | Estimate | ± s.e. | t-value | P |
|---|---|---|---|---|---|
| 8-isoprostane | Intercept | 3.13 | ± 0.13 | 23.60 | <0.001 |
| | Sex(males) | 0.11 | ± 0.13 | 0.82 | 0.413 |
| | Age(old) | 1.93 | ± 0.13 | 15.34 | <0.001 |
| | Sex x Age | -0.13 | ± 0.19 | -0.65 | 0.515 |
| o-tyrosine | Intercept | 3.28 | ± 0.15 | 22.15 | <0.001 |
| | Sex(males) | 0.17 | ± 0.14 | 1.21 | 0.229 |
| | Age(old) | 2.10 | ± 0.14 | 14.83 | <0.001 |
| | Sex x Age | -0.27 | ± 0.22 | -1.23 | 0.218 |
| 8-OHdG | Intercept | 2.37 | ± 0.11 | 20.87 | <0.001 |
| | Sex(males) | 0.16 | ± 0.12 | 1.34 | 0.182 |
| | Age(old) | 1.46 | ± 0.12 | 12.60 | <0.001 |
| | Sex x Age | -0.19 | ± 0.18 | -1.06 | 0.290 |
